# Evading non-essential fetal oocyte attrition maximizes the ovarian reserve

**DOI:** 10.1101/656645

**Authors:** Marla E. Tharp, Safia Malki, Alex Bortvin

## Abstract

Female reproductive success depends on the size and quality of a finite ovarian reserve. Paradoxically, mammals such as mice and humans eliminate up to 80% of the initial oocyte pool during development through the enigmatic process of fetal oocyte attrition (FOA). Here we report that the combined inhibition of retrotransposon L1 reverse transcriptase and the Chk2-dependent DNA damage checkpoint prevents FOA, thus preserving the entire fetal oocyte population. Remarkably, reverse transcriptase inhibitor AZT-treated *Chk2* mutant oocytes initially accumulate, but subsequently resolve L1-instigated genotoxic threats and differentiate, resulting in a maximized functional ovarian reserve. We conclude that FOA is a consequence of genotoxic stress that acts to preserve oocyte genome integrity, and is not an obligatory developmental program for oogenesis.

## Results and Discussion

The size and quality of the finite supply of immature oocytes established at birth are critical determinants of female reproductive lifespan and success (*1*). Paradoxically, the size of this starting ovarian reserve reflects only a minor share of the initial pool of fetal oocytes. In humans, as little as 20% of oocytes survive massive fetal oocyte attrition (FOA) to form the ovarian reserve at birth (*2*). Intriguingly, since the discovery of FOA in the 1960s, the rationale and mechanisms of this conserved process remain debated. Over the ensuing decades, inadequate oocyte nutrition, oocyte self-sacrifice, and meiotic DNA damage featured prominently in models attempting to explain this phenomenon (*3*). However, it remains unknown whether preventing FOA, if possible, would benefit or perturb fertility.

Previously, we implicated retrotransposon LINE-1 (L1), a highly abundant mobile element in mammalian genomes, in FOA in mice (Fig. 1A, B) (*4–6*). While DNA methylation and other mechanisms normally repress L1 (*7, 8*), epigenetic remodeling of primordial germ cells creates permissive conditions for L1 expression (*9*). In male germ cells, Piwi-interacting small RNAs (piRNAs) guide DNA methylation to rapidly repress transposons, whose excessive activity could otherwise lead to DNA damage, meiotic defects and germ cell death (*10–13*). In contrast, oocyte genomes remain demethylated throughout meiotic prophase I and until the onset of postnatal differentiation, expressing L1 during FOA (*14, 15*). In addition to correlating expression levels of L1-encoded ORF1p with DNA damage and oocyte fate, we also reported a remarkable protective effect of a reverse transcriptase inhibitor azidothymidine (AZT) on oocyte viability (*4, 16, 17*). Intriguingly, AZT treatment prevents oocyte death only temporarily, until embryonic day (E) 18.5. We surmised that the inability of AZT to prevent FOA completely is due to the persistent endonuclease activity of L1 ORF2p (*18*) resulting in excessive DNA damage and subsequent activation of the DNA damage checkpoint in E18.5 oocytes (*4, 19, 20*).

**Fig. 1.**
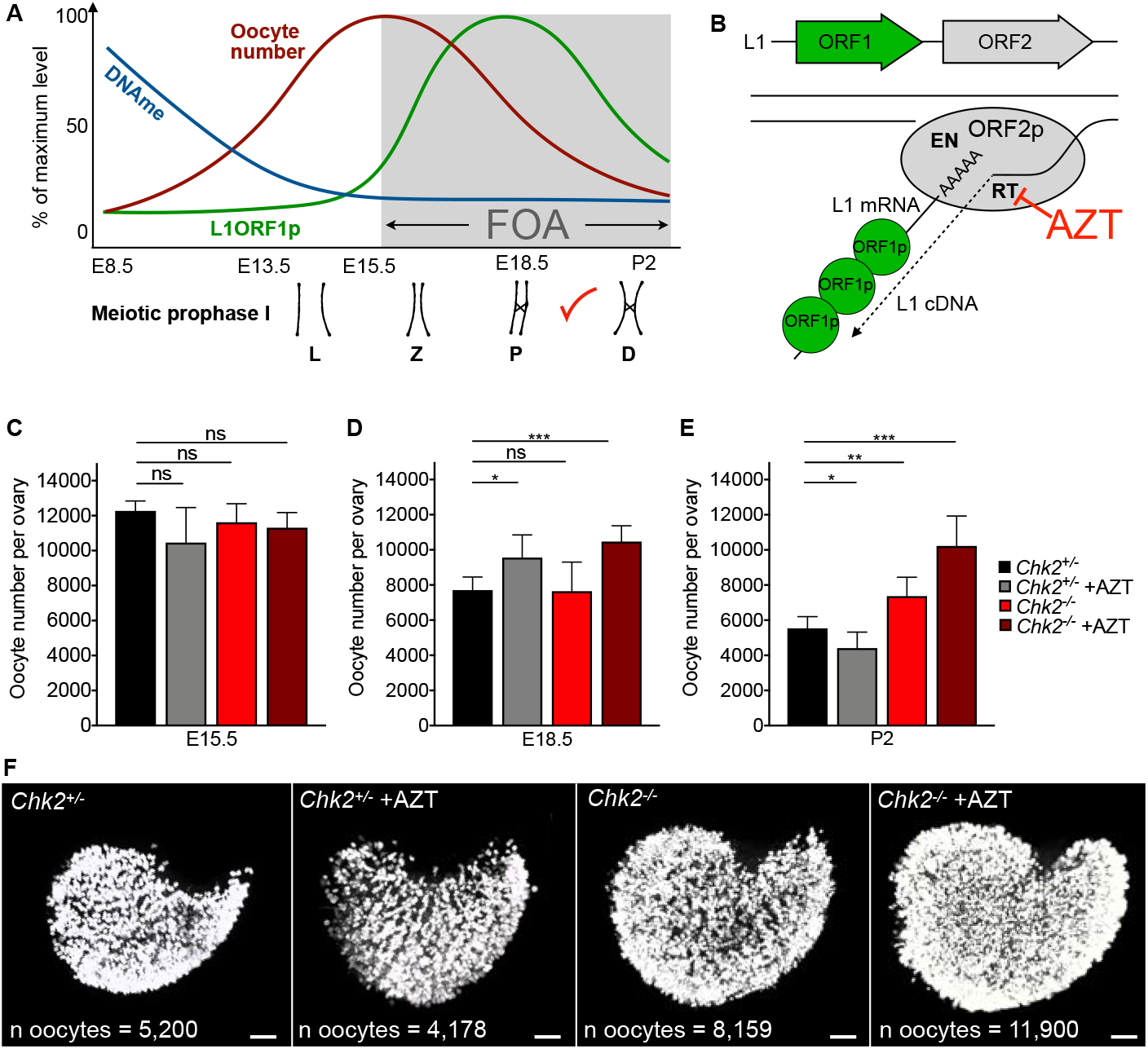
The dynamics and experimental evasion of Ll-mediated FOA. (**A**) Diagram of developmental programs concurrent with FOA and their dynamics, including DNA methylation (DNAme), L1 ORF1p expression, and meiotic prophase I (Leptotene (L), zygotene (Z), pachytene (P), and diplotene (D) stages indicated with red checkmark representing DNA damage checkpoint. (**B**) Diagram of L1 structure and retrotransposition mechanism. L1 propagation involves cytoplasmic packaging of L1 RNA in ribonucleoprotein particles along with L1-encoded RNA chaperon ORF1p and a reverse transcriptase (RT)/endonuclease (EN) ORF2p. Once in the nucleus, ORF2p endonuclease nicks DNA releasing a 3’-OH end to prime reverse transcription of L1 RNA into cDNA. The subsequent RNA removal from the L1 RNA:cDNAhybrid, second strand L1 DNA synthesis and ligation to genomic DNA produce new L1 insertions. AZT inhibits the RT activity of ORF2p. (**C-E**) Oocyte number per ovary at E15.5 (C), E18.5 (D) and P2 (E) in *Chk2*^+/−^ and *Chk2*^−/−^ untreated and AZT-treated ovaries. Mean + SD from at least three independent measurements shown. Statistics determined by unpaired t-test, ns p>0.05; *p<0.05; **p<0.01; ***p<0.001. (F) Whole-mount TRA98 labeling of P2 *Chk2*^+/−^ and *Chk2*^−/−^ untreated and AZT-treated ovaries. Oocyte number per ovary indicated on representative images. Scalebars represent 100μm.

To test if FOA is dependent on both L1 activity and L1-instigated activation of the DNA damage checkpoint, we first examined the role of checkpoint kinase 2 (CHK2), which transduces information regarding DNA double-stranded breaks (DSBs) to downstream apoptotic machinery in the mid-pachytene stage of meiotic prophase I (*21*). *Chk2* mutant mice are viable, fertile and, unlike in wild-type animals, unrepaired DSBs do not cause death of oocytes lacking CHK2 (*19*). However, whether CHK2 has a role in FOA under physiological conditions is unknown. We first compared the number of oocytes in *Chk2*^−/−^ to *Chk2*^+/−^ E15.5 ovaries at the onset of FOA. At this timepoint, there was no difference in oocyte number between *Chk2*^−/−^ and *Chk2*^+/−^ ovaries, suggesting that CHK2 does not impact primordial germ cell proliferation (Fig. 1C and fig. S1). We also analyzed the role of CHK2 in progression through meiotic prophase I based on synaptonemal complex morphology. We found no significant difference in the percent of total oocytes in each stage of meiotic prophase I between *Chk2*^−/−^ and *Chk2*^+/−^ ovaries at E15.5 (fig. S2A). However, in *Chk2*^−/−^ ovaries at E18.5, we observed an over-representation of diplotene stage oocytes, likely spared in the absence of DNA damage checkpoint activation in mid-pachynema (Fig. 1A and fig. S2B and table S1). We went on to compare oocyte number per ovary in *Chk2*^−/−^ and *Chk2*^+/−^ at E18.5 when approximately half of the oocytes are eliminated in wild-type mice, and at P2, the endpoint of FOA (*4*). At E18.5, the oocyte number is again comparable between *Chk2*^−/−^ and *Chk2*^+/−^ ovaries (Fig. 1D). However, by P2, *Chk2*^−/−^ ovaries contained significantly more oocytes than P2 *Chk2*^+/−^ ovaries, and a comparable number to E18.5 *Chk2*^−/−^ ovaries, suggesting no further FOA has occurred (Fig. 1E). These observations are consistent with the timing of DNA damage checkpoint activation in mid-pachynema as measured by upregulation of the transactivating p63 (TAp63) isoform that is downstream of CHK2 to trigger apoptosis at E18.5, but not at E15.5 (fig. S3). Therefore, CHK2 plays a role in FOA starting at E18.5, but not earlier.

Since the onset of CHK2 function in FOA at E18.5 coincided with the endpoint of the protective effect of AZT described previously (*4*), we tested if the combination of AZT treatment inhibition and DNA damage checkpoint would prevent FOA. We first validated the previous finding that daily administration of AZT to pregnant females starting at E13.5 prevents FOA between E15.5 and E18.5 but fails to maintain this effect until P2 in control *Chk2*^+/−^ mice (Fig. 1, C-E) (*4*). In contrast, AZT treatment of *Chk2*^−/−^ animals preserved more oocytes by P2 than either condition alone (Fig. 1E). In some cases, all fetal oocytes initially generated at E15.5 persist to P2 in *Chk2*^−/−^ +AZT conditions, resulting in a maximized oocyte supply (Fig. 1C-F). Notably, ending AZT treatment of *Chk2*^−/−^ mice at E18.5 did not maximize oocyte supply at P2 (fig. S4 and table S2). This evidence supports our hypothesis that FOA is a consequence of L1 reverse transcriptase activity throughout meiotic prophase I and activation of DNA damage checkpoint by L1 endonuclease activity and meiotic defects beginning in mid-pachynema.

To obtain a comprehensive picture of cell autonomous and cell non-autonomous mechanisms of oocyte loss during FOA and upon FOA evasion, we compared gene expression profiles of wild-type untreated and AZT-treated isolated oocytes and whole ovaries at the onset and at the peak of FOA (E15.5 and E18.5, respectively) (fig. S5, A - D). At E18.5, untreated samples contain surviving oocytes while AZT-treated samples contain rescued oocytes that should have been lost to FOA as well as surviving oocytes. Globally, RNA sequencing of ovaries and oocytes revealed similar gene expression profiles in both untreated and AZT-treated conditions at their respective developmental time points (fig. S6, A). This observation suggests that AZT treatment does not impact global gene expression significantly and that oocytes retain their cellular/developmental identities irrespective of their fate in FOA. However, closer analysis of gene expression between untreated and AZT-treated conditions at E18.5 shed light on differences related to FOA (fig. S6, B-E). Specifically, gene ontology enrichment analysis in E18.5 AZT-treated whole ovaries made apparent lower mRNA expression levels of a cohort of genes involved in the complement system of innate immunity compared to untreated ovaries (Fig. 2A-B). In contrast, E18.5 AZT-treated oocytes showed increased expression of the *CD55* gene also known as Decay Accelerating Factor DAF that protects cells from autologous complement attack (Fig. 2B). These observations indicated a role for the complement system in FOA that was further supported by the reduced occurrence of macrophages in ovaries that evaded FOA at E18.5 (fig. S7). Interestingly, antiviral innate immunity genes showed little ovarian and oocyte expression in all conditions, arguing against a prominent role of this pathway in FOA despite a strong response to L1 overexpression observed in other somatic tissues (Fig. 2B) (*22*).

**Fig. 2.**
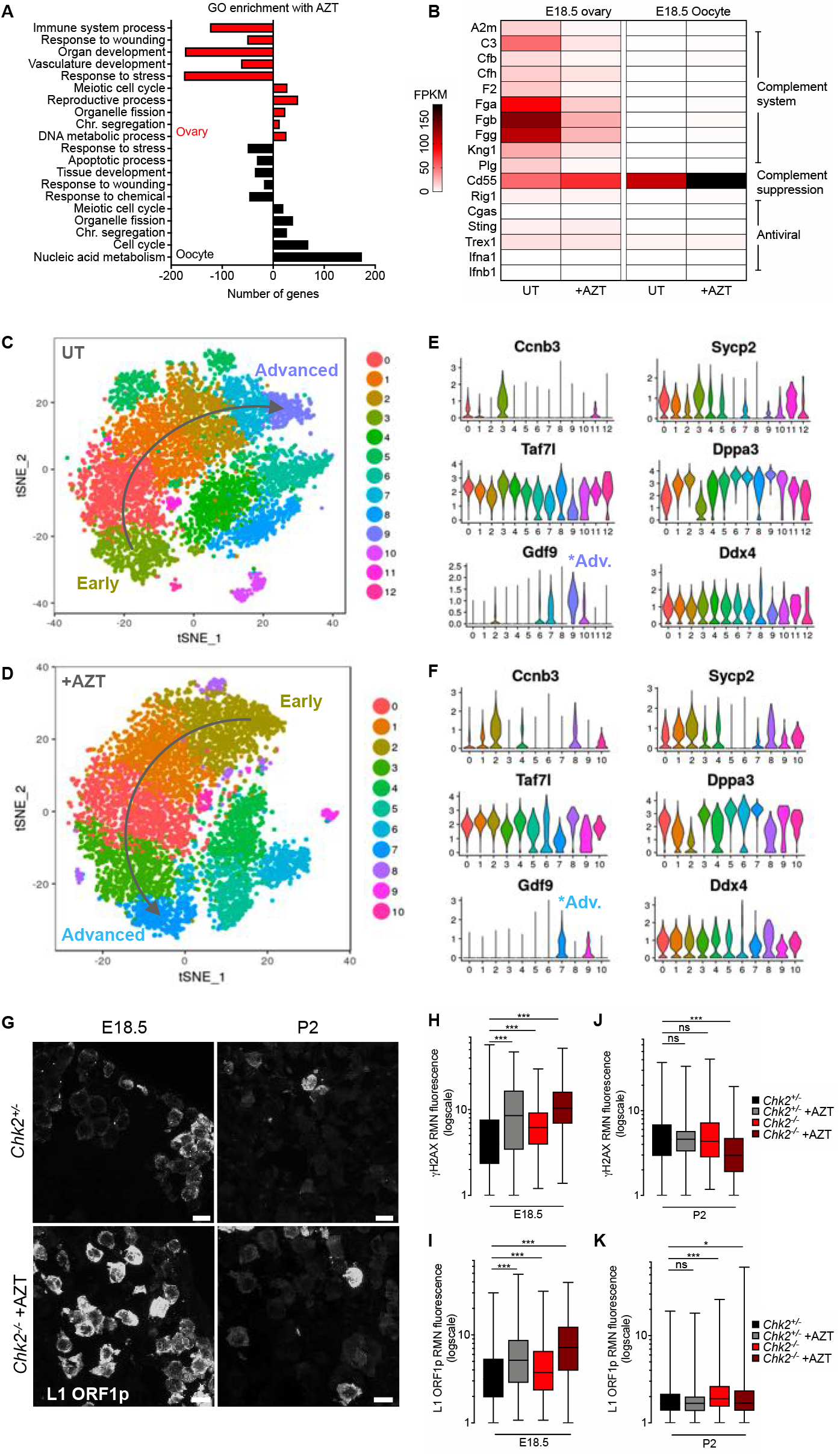
Gene expression analysis during FOA and upon FOA evasion. (**A**) Gene ontology enrichment upon AZT treatment in ovaries and oocytes. (**B**) Differential gene expression related to the innate immune system between untreated (UT) and AZT-treated conditions in E18.5 ovaries and oocytes. (**C-D**) Re-clustering of oocyte-specific clusters in E18.5 untreated (**C**) and AZT-treated (**D**) ovaries by t-SNE. Early to advanced oocyte developmental trajectories indicated by arrows. (**E-F**) Expression of early, intermediate and advanced genes in untreated (**E**) and AZT-treated (**F**) oocyte clusters. *Ccnb3* is expressed early, *Sycp2, Taf7l* and *Dppa3* are expressed in intermediate stages, and *Gdf9* is expressed in advanced oocytes. *Ddx4* is expressed in all oocytes. (**G**) Immunofluorescence labeling of L1 ORF1p in *Chk2*^+/−^ and *Chk2*^−/−^ +AZT ovaries at E18.5 and P2. Scalebar indicates 10μm. (**H-I**) Box and whiskers plot of RMN γH2AX levels (**H**) and L1 ORF1p levels (**I**) in *Chk2*^+/−^ and *Chk2*^−/−^ untreated and AZT-treated oocytes at E18.5. (**J-K**) Box and whiskers plot of RMN γH2AX levels (**J**) and L1 ORF1p levels (K) in *Chk2*^+/−^ and *Chk2*^−/−^ untreated and AZT-treated oocytes at P2. Statistics determined by Kolmogorov-Smirnov test, ns p> 0.05; *p<0.05; ***p<0.001.

L1 expression remains a prominent feature distinguishing AZT-treated from untreated oocytes during FOA and dictates the fate of an individual oocyte (fig. S8) (*4*). To better understand transcriptional heterogeneity between fetal oocytes and gene expression profiles related to FOA, we performed single-cell RNA sequencing of E18.5 ovaries in untreated and AZT-treated conditions. Cells from both wild-type untreated and AZT-treated E18.5 ovaries were clustered and re-clustered into oocyte-specific populations (Fig. 2, C - D and fig. S9, A - D). Oocytes separated into early, intermediate, and advanced developmental stages relative to the E18.5 timepoint based on expression of established germ-cell markers (Fig.2, E-F and fig. S9, E-F, and table S4). Consistent with the earlier results of bulk oocyte RNA sequencing, AZT treatment did not change the E18.5 oocyte transcriptome significantly, nor the separation into early, intermediate, and advanced developmental stages. However, the proportion of oocytes present at each stage differed slightly. Consistent with the predominant activity of FOA in early meiotic prophase I between E15.5 and E18.5, and increased expression of meiotic genes in bulk AZT-treated oocytes, we observed a greater proportion of oocytes cluster into the early and intermediate developmental stages in the AZT-treated sample, while the untreated sample contained more oocytes expressing advanced markers such as *Gdf9*(Fig. 2E-F). Cumulatively, gene expression analyses suggest that oocytes surviving and perishing in FOA are fundamentally similar, with an over-representation of early and intermediate stage oocytes in conditions of FOA evasion, which is consistent with the predominant oocyte loss in early meiotic prophase I.

To determine how oocytes evading FOA progress through development in the presence of substantial genotoxic stress, we measured the nuclear abundance of γH2AX (a DSB marker) and L1 ORF1p in individual oocytes of *Chk2*^+/−^, *Chk2*^−/−^, and AZT-treated *Chk2*^+/−^ and *Chk2*^−/−^ ovaries. Consistent with previous findings (*4*), a population of AZT treated E18.5 *Chk2*^+/−^ or *Chk2*^−/−^ oocytes showed higher γH2AX abundance compared to untreated ovaries (Fig. 2H). Additionally, AZT-treated *Chk2*^+/−^ and *Chk2*^−/−^ E18.5 ovaries contained a population of oocytes with high levels of L1 ORF1p and L1 ORF1 mRNA never observed in untreated controls (Fig. 2G, I and fig. S8). Although untreated *Chk2*^−/−^ showed higher L1 ORF1p levels compared to controls in E18.5 oocytes, AZT treatment increased L1 ORF1p levels further (Fig. 2I). Thus, inhibition of both L1 and the DNA damage checkpoint results in the appearance of E18.5 oocytes exhibiting L1 overexpression and excessive genome damage. Unexpectedly, although *Chk2*^−/−^ and *Chk2*^−/−^ +AZT P2 oocytes still showed elevated L1 ORF1p levels compared to *Chk2*^+/−^ and *Chk2*^+/−^ +AZT oocytes, P2 oocytes in all conditions showed significantly reduced L1 ORF1p and γH2AX levels compared to E18.5 (Fig. 2, G, J-K and table S2E-H). An FOA-independent reduction of L1 expression and γH2AX levels indicates that while FOA acts as quality control for genome integrity, given a chance, an oocyte can ultimately reduce genotoxic threats.

While previous studies showed significant DNA damage repair capacity of *Chk2*^−/−^ mutant postnatal oocytes (*19*), the unexpected reduction of L1 ORF1p levels prompted us to examine the involvement of piRNAs in FOA. Indeed, robust downregulation of L1 ORF1p expression in E18.5-P2 oocytes coincided with increased expression of piRNA pathway genes including *Mili* that encodes the predominant Piwi family protein in wild-type fetal oocytes (Fig. 3, A-B) (*23*). Consistent with piRNA pathway activation, RNAs of 26-27 nucleotides in length (characteristic of piRNA) became highly abundant in E18.5 and P2 wild-type ovaries and superseded RNAs of 22-23 nucleotides (characteristic of endo-siRNAs) dominating the E15.5 RNA profile (Fig. 3C). Mapping of small RNA reads to repetitive elements revealed a massive increase in antisense small RNAs targeting evolutionarily young L1MdA and L1MdT elements at E18.5 and P2 compared to E15.5 (Fig. 3D-E and table S5). Importantly, *Mili* mutant P2 ovaries had negligible amounts of piRNAs based on the absence of the 26-27 nucleotide long small RNA population, proving that piRNA production at this time is Mili-dependent (Fig. 3F). However, we found that P2 oocytes evading FOA in *Mili*^−/−^;*Chk2*^−/−^ double-mutant mice treated with AZT still downregulated L1 ORF1p levels (Fig. 3, G and H and table S6). Cumulatively, these results suggest that oocytes use multiple mechanisms to reduce L1 upon evading FOA, one candidate being HSP90a, expression of which negatively correlates with L1 ORF1p in human germ cells and is implicated in post-transcriptional transposon repression in mice (*24, 25*). Indeed, we found that *Hsp90a* family genes are significantly expressed and upregulated between E15.5 and E18.5 oocytes (Fig 3A).

**Fig. 3.**
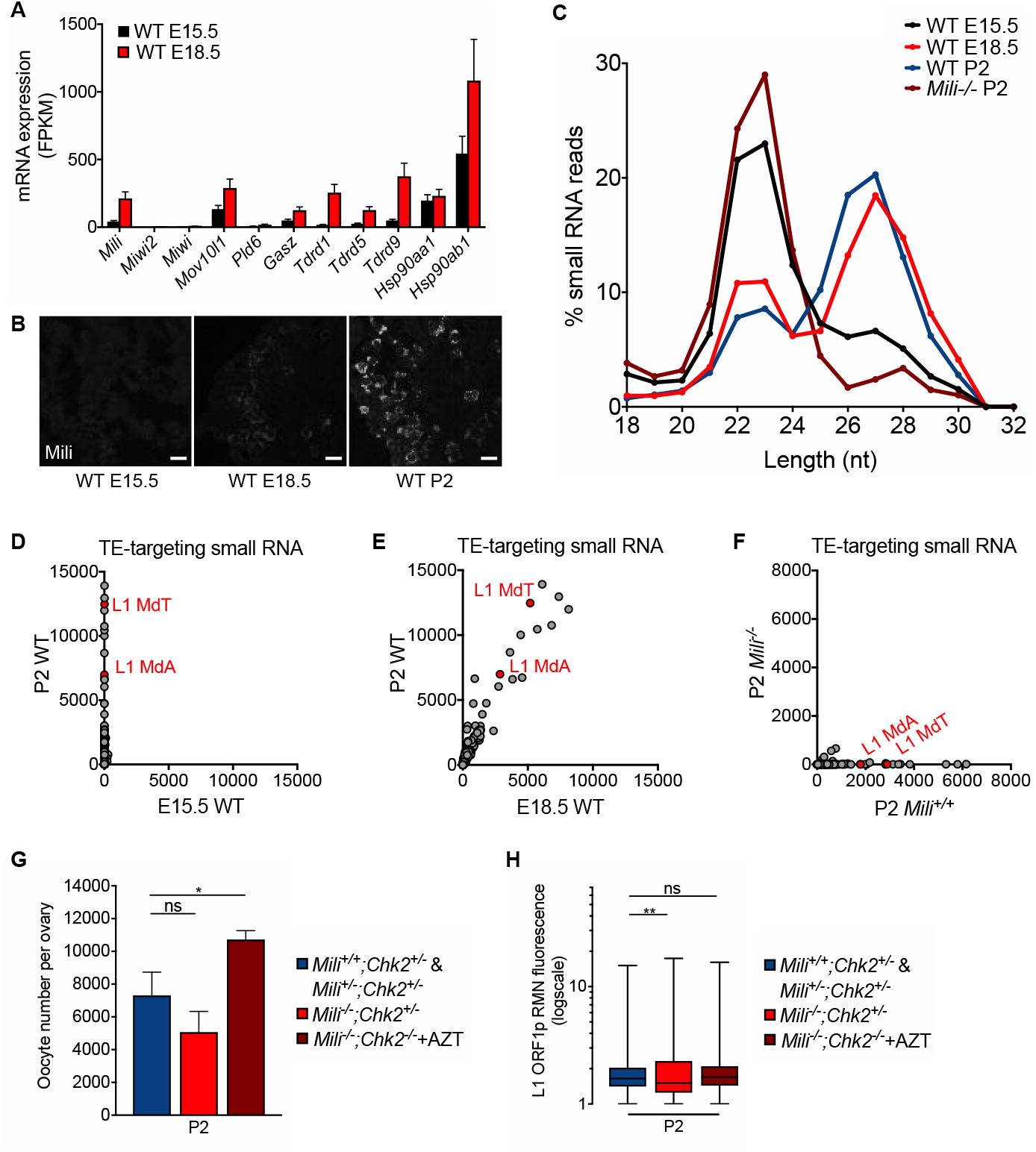
Downregulation of L1 and DNA damage repair in oocytes evading FOA. (**A**) mRNA expression of piRNA biogenesis and Hsp90a family genes in E15.5 and E18.5 WT oocytes. (**B**) Mili labeling of E15.5, E18.5 and P2 WT ovary sections. Scalebars represent 15μm. (**C**) Size distribution of 18-32 nt reads in small RNA libraries from E15.5, E18.5, and P2 WT and P2 *Mili*^−/−^ ovaries. (**D-F**) Abundance of antisense transposon-targeting small RNAs in E15.5 vs P2 WT, E18.5 vs P2 WT (E), and P2 *Mili*^+/+^ vs P2 *Mili*^−/−^ (F) ovaries. (**G**) Oocyte number in *Mili;Chk2* double mutant and control P2 ovaries. Statistics determined by unpaired t-test, ns p>0.05; *p<0.05. (**H**) Box and whiskers plot of RMN L1ORF1p levels in *Mili;Chk2* double mutant and control ovaries. Statistics determined by Kolmogorov-Smirnov test, ns p> 0.05; **p<0.01.

The striking reduction of L1 expression and repair of DNA breaks in oocytes evading FOA prompted us to test their differentiation and developmental capacity. To assess oocyte differentiation, we analyzed the Balbiani body, an organelle aggregate that contains the Golgi complex, mitochondria, and RNAs, and whose size increases with oocyte growth and differentiation (*26*). We quantified Balbiani body size at E18.5 and P2, based on Golgi content area. We found an increase in Golgi content from 40μm^2^ to 70μm^2^ in *Chk2*^+/−^ untreated control oocytes (Fig. 4B). Importantly, oocytes in *Chk2*^−/−^ untreated and *Chk2*^+/−^ and *Chk2*^−/−^ AZT-treated conditions also experienced a significant increase in Golgi content (Fig. 4A). Therefore, oocytes allowed to reach the P2 stage, regardless of L1 expression or amount of DNA damage initially accumulated at E18.5, have the potential to differentiate if FOA is prevented (Fig. 4B).

**Fig. 4.**
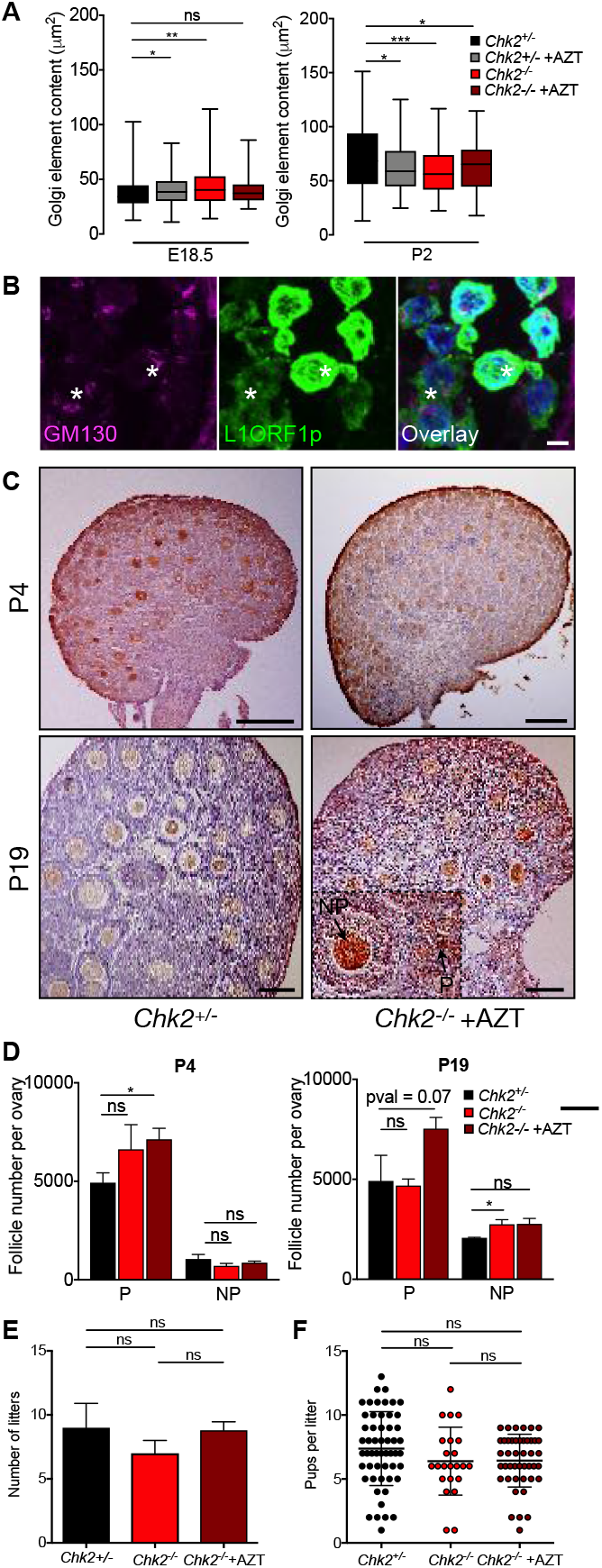
Normal oogenesis and fertility in mice evading FOA. (**A**) Box and whiskers plots of Golgi element content area (μm^2^) in *Chk2*^+/−^ and *Chk2*^−/−^ untreated and AZT-treated ovaries at E18.5 and P2. Statistics determined by Kolmogorov-Smirnov test, ns p> 0.05; *p<0.05; **p<0.01; ***p<0.001. (**B**) GM130 and L1 ORF1p labeling in *Chk2*^−/−^ +AZT ovaries at E18.5. White asterisks indicate oocytes of high and low L1 ORF1p expression with comparable Balbiani body sizes. Scalebars represent 5μm. (**C**) MVH labeling and hematoxylin counter-staining of P4 and P19 *Chk2*^+/−^ and *Chk2*^−/−^ +AZT ovaries. Scalebars represent 100μm. Inset shows example of primordial (P) and non-primordial (NP) follicles. (**D**) Quantification of P and NP follicles in P4 and P19 ovaries. Statistics determined by unpaired t-test, ns p>0.05; *p<0.05. (**E**) Number of litters per *Chk2*^+/−^, *Chk2*^−/−^, and *Chk2*^−/−^ +AZT female crossed to *Chk2*^+/−^ male over the course of 10 months. Only females that survived the duration of the assay were counted. Statistics determined by unpaired t-test, ns p>0.05. **(F)** Pups per litter from *Chk2*^+/−^, *Chk2*^−/−^, and *Chk2*^−/−^ +AZT females crossed to *Chk2*^+/−^ males. Statistics determined by unpaired t-test, ns p>0.05.

To assess the significance of FOA for folliculogenesis, we quantified growing follicle types in P4 and P19 ovaries of *Chk2*^+/−^, *Chk2*^−/−^, and *Chk2*^−/−^ +AZT females. P4 is an early stage of folliculogenesis at which the transition from primordial to primary follicles occurs through encapsulation of oocytes with somatic granulosa cells. P19 stage ovaries have further progressed through folliculogenesis and contain all follicle types that form by further accumulation of granulosa cell layers and theca cells around oocytes, but before ovulation. We observed the characteristic growing follicle types at P4 and P19 in both *Chk2*^−/−^ +AZT and control ovaries, suggesting that folliculogenesis is not impeded in the absence of FOA (Fig. 4, C - D). Quantification of primordial and non-primordial follicles in P4 ovaries that have evaded FOA revealed an increased number of primordial follicles compared to controls and comparable numbers of non-primordial follicles (Fig. 4, C - D). P19 ovaries that do not undergo FOA also contain more follicles, but the increase observed was less significant (Fig. 4, C - D).

To test the consequences of possessing increased numbers of primordial follicles on fertility as a result of FOA prevention, we crossed females born to *Chk2*^−/−^ +AZT-treated females that have reached reproductive maturity to *Chk2*^+/−^ males and monitored the number of litters and litter size compared to untreated *Chk2*^−/−^ and *Chk2*^+/−^ females. Both litter number and litter size over the course of 10 months was comparable between the three conditions, suggesting that FOA is not obligatory for fertility, nor is it detrimental to have an increased number of primordial follicles in the ovarian reserve (Fig. 4F and table S7).

Cumulatively, this work identifies FOA as a consequence of genotoxic stress during fetal oogenesis that is triggered by excessive L1 reverse transcriptase and endonuclease activities leading to DNA damage and meiotic defects. We showed that oocytes can be manipulated to evade FOA, downregulate L1 expression, and maximize the functional ovarian reserve without disrupting oogenesis or reducing fertility. Therefore, in mammals, massive oocyte loss is not an obligatory developmental program for oogenesis. Crucially, due to the conservation of FOA and L1 expression in human oocytes, this work is of biomedical relevance for human female fertility as it identifies an avenue for the improvement of the ovarian reserve by inhibiting L1 activity.

## Acknowledgments

We thank F. Tan and A. Pinder for assistance with RNA sequencing and data analyses; R. Juste and C. Hussey for technical assistance; E. Dikovsky for assistance with animal care; Tak W. Mak for providing *Chk2* mutant mice, H. Kazazian for providing *Mili* mutant mice; Z. Zhang for assistance and reagents with small RNA library preparation; and Mark Van Doren for critical reading of manuscript.

## Funding

This work was supported by NIH grants R21 HD090514 to A.B. and F31 HD088053 to M.E.T.

## Author contributions

Conceptualization - A.B., M.E.T., S.M.. Funding acquisition - A.B., M.E.T. Investigation - M.E.T, S.M.. Supervision - A.B. Writing original draft, review and editing - M.E.T., A.B.

## Data and materials availability

The NCBI Sequence Read Archive project number for all high-throughput sequencing data is: PRJNA543598.

## Supplementary Materials

Materials and Methods

Tables S1 – S7

Fig S1 – S9

References (27 - 35)

## Supplementary Materials

### Materials and Methods

#### Animals

For this study, *Chk2*^−/−^ mice in a mixed C57B1/6 and 129X1/Sv genetic background were used (*27*). *Chk2*^−/−^ mice were backcrossed one time to C57B1/6 to generate *Chk2*^+/−^ controls. We chose *Chk2*^+/−^ as a control to account for the mixed genetic background resulting in a significant increase in oocyte number compared to wild-type mice of pure C57B1/6 background (fig. S1). To determine how genetic background contributed to differences in oocyte number between *Chk2*^−/−^ and C57B1/6 wild-type mice, we backcrossed *Chk2*^−/−^ for 4 generations. This reduced the percent genome containing homozygous 129X1/Sv SNPs from approximately 4% to 0.03% (SNP genotyping by DartMouse, Dartmouth School of Medicine). *Mili*^+/+^, *Mili*^+/−^, and *Mili*^−/−^ mice used were in C57B1/6 genetic background. *Mili*^−/−^;*Chk2*^−/−^ mice and control littermates were generated by crossing *Chk2*^−/−^ and *Mili*^+/−^ animals. Wild-type mice of CD1 (Charles River Laboratories) genetic background were used for all quantitative PCR, mRNA and small RNA sequencing experiments unless otherwise noted. All experimental procedures were performed in compliance with ethical regulations and approved by the IACUC of the Carnegie Institution for Science.

#### AZT treatment

50mg/kg/day AZT was administered daily by gavage to ~8-month-old pregnant female from E13.5 until experiment end point. AZT (Sigma Aldrich, Cat# A2169) was diluted to 5mg/ml in nuclease free water, aliquoted and stored at −20°C.

#### Oocyte and follicle quantification

We quantified oocytes per ovary at E15.5, E18.5, and P2 using a whole-mount immunofluorescence and tissue clearing method previously described (*28*). Ovaries were dissected and labeled with the germ cell-specific antibody TRA98 (*29*). After immunofluorescence, samples were treated with ScaleA2 clearing reagent for 7 days, changing solution each day. Confocal imaging through the entire tissue using SP5 confocal microscope (Leica) followed by 3D reconstruction of Z-stacked images and spot and surface analysis using Imaris software (Bitplane) were performed. For statistical analysis, average number of oocytes per ovary was counted in embryos from at least 3 different litters (table S2A). Variability between numbers due to timing of plug during the day as well as natural variation between embryos that is more apparent at earlier stages between litters and embryos. Statistical significance was determined using two-tailed unpaired Student’s t-test (table S2B). In the case of *Mili;Chk2* double mutants, we quantified oocytes labeled with germ cell-specific antibody TRA98 and nuclei counterstained with DAPI by scoring every 5^th^ section through the entire ovary, and estimating total number per ovary. Statistical significance was determined using two-tailed unpaired Student’s t-test (table S6A-B).

We quantified primordial and non-primordial (primary, secondary, antral) follicles at P4 and P19 by performing immunohistochemistry and DAB staining. We quantified follicles in every 5^th^ section through the entire ovary, then estimated total number per ovary. Sections were labeled with the cytoplasmic germ cell marker MVH and nuclei counterstained with hematoxylin. MVH-positive follicles were quantified and categorized as primordial or non-primordial based on the number of somatic cell layers surrounding the oocyte. At least three ovaries from three different females were quantified for each experimental group and two-tailed unpaired Student’s t-test used to determine statistical significance (table S7A-B).

#### Immunostaining

Whole-mount immunofluorescence and immunofluorescence protocols on cryosections and meiotic spreads were performed as previously described (*4, 28*). For immunohistochemistry with DAB staining, ovaries were fixed in Bouin’s overnight at 4°C, transferred to 70% ethanol overnight, and embedded in paraffin. Ovaries were sectioned into 10μm slices. Sections were deparaffinized by washing slides in Citrisolv (Decon Labs) 3x 15 minutes, re-hydrated through graded ethanol washes, blocked with hydrogen peroxide for 10 minutes, avidin block for 15 minutes (Vector Laboratories), biotin block (Vector Laboratories), and goat serum block (Vector Labs cat # PK-4001). Samples were incubated overnight at 4°C with anti-DDX4/MVH and for 30 minutes at room temperature with biotinylated goat anti-rabbit IgG secondary antibody (Vector Labs cat # PK-4001). Samples were incubated 30 minutes at room temperature with Vectastain ABC reagent (Vector Labs cat # PK-4001) followed by DAB detection. Slides were dipped in hematoxylin, rehydrated in ethanol, dipped in Citrisolv and mounted.

#### Microscopy

Imaging of whole-mount ovaries and ovary sections was performed using TCS-SP5 laser-scanning confocal microscope (Leica, Buffalo Grove, IL), histological sections using Nikon Eclipse E800 microscope equipped with a Diagnostic Instruments model 2.3.1 digital camera, and meiotic spreads using Olympus BX61 microscope equipped with a Hamamatsu C4742-95 digital Camera as previously described (*4, 28*). Image analysis was completed using Imaris (Bitplane) and ImageJ.

#### Antibodies

The following primary antibodies and dilutions were used:

Anti-Germ cell-specific antigen antibody [TRA98], rat monoclonal, abcam cat. # ab82527, diluted to 1:500 for immunofluorescence.

Anti-L1 ORF1p (full length protein), rabbit polyclonal, diluted to 1:500 for immunofluorescence (*4*).

Anti-GM130, mouse monoclonal, BD Biosciences cat. # 610822, diluted to 1:200 for immunofluorescence.

Anti-phospho-Histone H2A.X (Ser139) clone JBW301, mouse monoclonal, Millipore Sigma cat. # 05-636, diluted to 1:1000 for immunofluorescence.

Anti-SYCP3, rabbit polyclonal, abcam cat. # ab15093, diluted to 1:500 for immunofluorescence.

Anti-PIWIL2, rabbit polyclonal, abcam cat. # ab181340, diluted to 1:50 for immunofluorescence.

Anti-F4/80 antibody [CI:A3-1], rat monoclonal, abcam cat. # ab6640, diluted to 1:100 for immunofluorescence.

Anti-DDX4/MVH, rabbit polyclonal, abcam cat. # ab13840, diluted to 1:200 for immunostaining on paraffin sections and 1:1000 for western blot.

Anti-p63(4A4), mouse monoclonal, Santa Cruz Biotechnology cat. # sc-8431, diluted to 1:500 for western blot.

The following secondary antibodies and dilutions were used:

Alexa donkey ant-rabbit 488 (Invitrogen, cat # A-21206) diluted 1:1000 for immunofluorescence.

Alexa donkey anti-rabbit 568 (Invitrogen, cat # A10042) diluted 1:1000 for immunofluorescence.

Alexa donkey anti-mouse 488 (Invitrogen, cat # A-21202) diluted 1:1000 for immunofluorescence.

Alexa donkey anti-mouse 594 (Invitrogen, cat # A-21203) diluted 1:1000 for immunofluorescence.

Alexa donkey anti-rat 647 (Invitrogen, cat # 150155) diluted 1:1000 for immunofluorescence. Goat anti-mouse IgG (H+L)-HRP Conjugate (BioRad cat # 1721011) diluted 1:2000 for Western blot.

Goat anti-rabbit IgG (H+L)-HRP Conjugate (BioRad cat # 1721019) diluted 1:2000 for Western blot.

#### Real-time RT-PCR

We isolated RNA from FACS-sorted oocytes and somatic cells using TRIZOL reagent (Invitrogen). RNA was DNase treated using TURBO DNA-free kit (Ambion). cDNA synthesis reactions were performed using oligo dT and Superscript III First-Strand Synthesis System for RT-PCR (Invitrogen). cDNA was diluted equally and added to the qPCR reactions containing SsoAdvanced Universal SYBR Green Supermix (Bio-Rad). RT-PCR was performed on CFX96 Touch Real-Time PCR Detection System to detect SYBR Green. Relative quantities were analyzed using AACt methods with Actb as the housekeeping control gene. Primer sequences used for RT-PCR experiments include F-actb: CGG TTC CGA TGC CCT GAG GCT CTT; R-actb: CGT CAC ACT TCA TGA TGG AAT TGA; F-mvh: TGG CAG AGC GAT TTC TTT TT; R-mvh: CGC TGT ATT CAA CGT GTG CT; F-L1ORF1: ATG GCG AAA GGT AAA CGG AG; R-L1ORF1: AGT CCT TCT TGA TGT CCT CT.

#### Western blot

6-12 whole ovaries were lysed in RIPA buffer containing 50mM Tris-HCl pH=8, 150mM NaCl, 1%NP40, 1%SDS, 1mMEDTA, and 10% glycerol. 1mM PMSF and 1mM Halt protease and phosphatase inhibitor cocktail were added to buffer just before lysis. Ovaries were homogenized using RNase-free pestle and protein quantified using BCA. Lysates were run on 12% polyacrylamide running, 4% polyacrylamide stacking gel. Proteins transferred overnight at 4°C to PVDF membrane that had been activated for 15 seconds in 100% methanol followed by 2 minutes water and 15 min in transfer buffer. Membrane rinsed with PBS + 0.05% Tween-20 and blocked with 5% nonfat milk in PBS + 0.05% Tween-20 for 1 hour at room temperature. Primary antibodies were incubated overnight at 4°C in blocking buffer. Secondary antibodies used at 1:2000 and incubated 1 hour at room temperature. Detection by ECL was performed.

#### Meiotic chromatin spread preparation

Ovaries were dissociated into a single-cell suspension using dissociation buffer containing 0.025% trypsin, 2.5 mg/mL collagenase, and 0.1mg/mL DNase I. One volume of Hypotonic buffer (30mM Tris-HCl, pH 8.2, 50mM sucrose, 17mM sodium citrate) was added to the cell suspension and set on nutator for 30 minutes. Cells were pelleted and supernatant replaced with 100μM sucrose, pH 8.2 solution. Approximately 600μl sucrose solution per ovary pair. Slides were dipped in fixative (1% PFA, 0.15% Triton X-100, pH 9.2) and 20μL resuspended cells pipetted along bottom edge. Cells were slowly spread around slide by tilting the slide gently. Slides were dried in humid chamber for 2 hours, then treated with 0.08% Photo-Flo (Kodak). Slides used immediately for immunostaining or stored at −80°C.

#### Analysis of L1ORF1p and gH2AX nuclear fluorescence

Ovary sections of 8μm thickness were stained with DAPI, germ cell marker TRA98, and L1 ORF1p. Confocal stacks were taken through the section. Imaris bitplane was used to generate a surface around each DAPI-positive nucleus in TRA98-positive germ cells. Then, relative mean nuclear (RMN) fluorescence was calculated for the channel containing L1 ORF1p signal within the surface. This procedure was used in the same manner to calculate gH2AX RMN fluorescence. Each germ cell RMN value was then divided by the average of three RMN values from TRA98-negative somatic cell nuclei that should not contain L1 ORF1p nor gH2AX to normalize for background fluorescence. On average, about 200 oocytes were quantified per experimental group. Oocytes come from at least 3 different ovaries and 2 different litters unless noted otherwise (table S2C, E and table S6C). Statistical significance was determined using two-tailed Kolmogorov-Smirnov test (table S2D, F and table S6D).

#### Analysis of Golgi element area

Ovary sections of 8μm thickness were stained with DAPI, germ cell marker TRA98 and GM130 (*26*). Confocal stacks were taken through the section. Imaris bitplane was used to generate a surface around each GM130-positive Balbiani body region in a TRA98-positive cell. Area of surface generated for GM130 channel was calculated. Each bar represents 70 to 200 individual oocytes measured. Oocytes come from at least 3 different ovaries and 2 different litters unless noted otherwise (table S2G). Statistical significance was determined using two-tailed Kolmogorov-Smirnov test (table S2H).

#### FACS sorting

Ovaries were dissociated into a single-cell suspension using dissociation buffer containing 0.025% trypsin, 2.5 mg/mL collagenase, and 0.1mg/mL DNase I. Cell suspensions were filtered using 40μm filter. Oocytes were FACS-sorted from remaining ovarian somatic cells based on size and complexity using forward and side scatter parameters (fig. S6A-C) (*30*). Propidium iodide treatment for 10 minutes and gating for negative red fluorescence was used to eliminate dead cells from the sort. BD FACS Aria III sorter was used.

#### Bulk RNA-Seq analysis

RNA was extracted using Trizol reagent (Invitrogen), DNaseI-treated using TURBO DNA-free kit (Ambion), and libraries generated using ribo-zero kit. 75bp unpaired, single-end reads were sequenced on Illumina Next-seq 500 system. Remaining rRNA sequences were removed computationally using bowtie by aligning reads to mm10 rRNA genome. Reads that did not align were subsequently mapped to mm10 genome using Tophat splice aligner (*31*). To determine differential gene expression, cuffdiff was used followed by cummeRbund in R to obtain FPKM values and generate plots (fig. S7A-B and table S3A-D). GO pathway enrichment analyses was performed using DAVID Bioinformatics Resources 6.8 (table S3E-H) (*32, 33*).

#### Single-cell RNA-Seq analysis

Ovaries were dissociated into a single-cell suspension using dissociation buffer containing 0.025% trypsin, 2.5 mg/mL collagenase, and 0.1mg/mL DNase I. Cells were pelleted and washed with PBS 3 times. Viability and cell count were determined using trypan blue staining and Countess II automated cell counter. Samples with greater than 90% viability and 1 million cells per milliliter were used for sequencing. GEM generation, barcoding, and library construction were performed with 10x genomics Chromium Genome Reagent Kit (v2 Chemistry). Libraries were sequenced on Illumina Next-Seq 500 system. For the untreated sample, 16,448 cells with 48,924 mean reads per cell were sequenced. For the AZT-treated sample, 15,551 cells with 48,450 mean reads per cell were sequenced. Differential gene expression and clustering analysis was performed using Cell Ranger 3.0 and Seurat packages (table S4A-D) (*34*).

#### Small RNA-Seq analysis

Total RNA was extracted from whole ovaries using mirVana miRNA isolation kit. RNA was run on 15% urea gel and 18-35 nucleotide region excised. Small RNAs were eluded from gel slice with 0.3M NaCl overnight at room temperature. 3’ and 5’ adapters were added and reverse transcription reaction performed to generate cDNA. Libraries were PCR amplified and 75 or 150 bp reads sequenced on Illumina Next-Seq 500 system. piPipes small RNA analysis pipeline was used to determine small RNA length and align reads to repeats (table S5A-D) (*35*) (Han et al, 2015). For small RNA length distribution, rRNA and miRNA hairpins were removed. For repeat alignment and quantification, reads were normalized to miRNA. At least three pairs of ovaries were used per sample. WT samples were of CD1 genetic background. *Mili*^+/+^ and *Mili*^−/−^ were of B6 genetic background.

#### Fertility assay

*Chk2*^−/−^ females that were treated with AZT during their fetal development and raised to adults were crossed to *Chk2*^+/−^ males. *Chk2*^−/−^ females that were untreated as well as *Chk2*^+/−^ females that were untreated were crossed to *Chk2*^+/−^ males for controls. The number of live pups per litter at the day of birth from 6 *Chk2*^−/−^ +AZT females, 3 *Chk2*^−/−^ females, and 6 *Chk2*^+/−^ females were monitored for at least 10 months (table S7C). Number of litters over 10 months were reported in females that survived the duration of the assay. Statistical significance was determined using unpaired Student’s t-test (table S7).

**Fig. S1.**
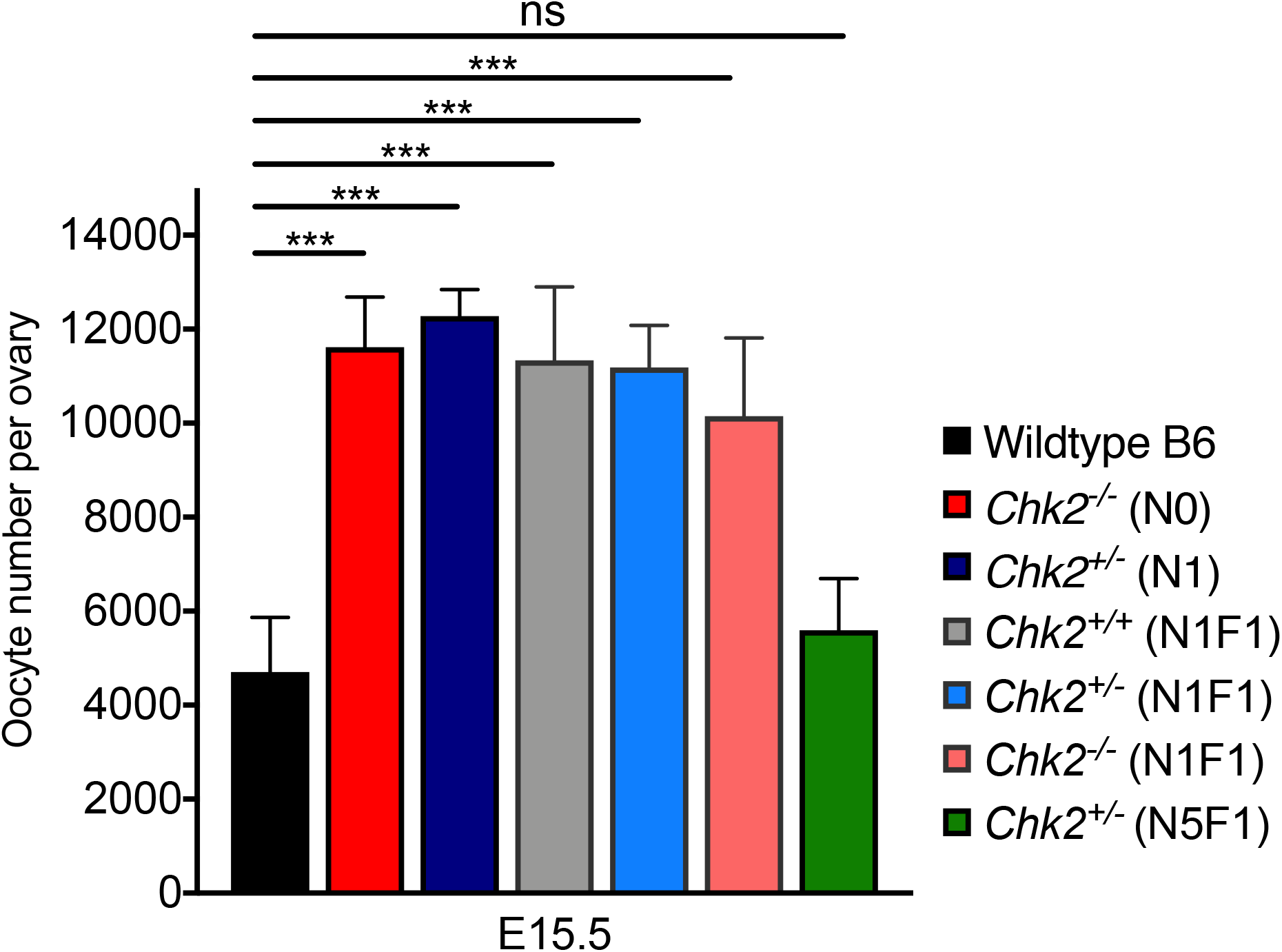
Contribution of genetic background to maximum oocyte number. Quantification of ovaries at E15.5 in *Chk2*^−/−^ of mixed C57B1/6 and 129X1/Sv genetic background and *Chk2*^−/−^ backcrossed to C57B1/6 to remove homozygous 129X1/Sv genome content (table S2C-D). N1, N1F1, and N5F1 backcrosses were compared. Oocyte number decreases with decreasing 129X1/Sv genome content, and N5F1 embryos showed oocyte number per ovary that is comparable to pure C57B1/6 embryos. Differences in oocyte number at E15.5 with genetic background are independent of the *Chk2* mutation. *Chk2*^+/−^ (N1) and *Chk2*^−/−^ (N0) results from Fig. 1C and table S2A-B. Statistical significance was determined by unpaired t-test, ns p>0.05; ***p<0.001.

**Fig. S2.**
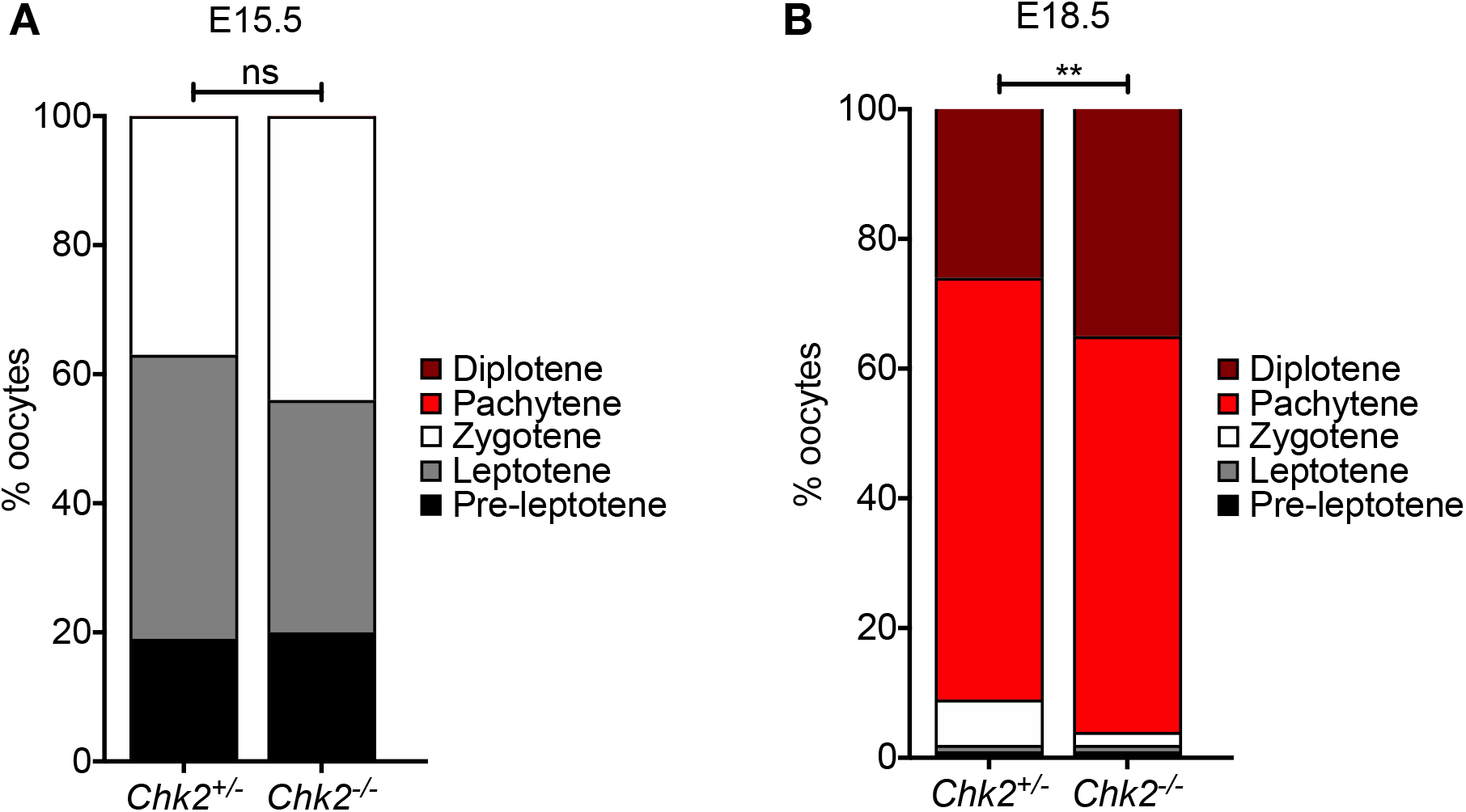
The role of Chk2 in meiotic progression. (**A-B**) Comparison of meiotic progression between *Chk2*^+/−^ and *Chk2*^−/−^ ovaries at E15.5 (**A**) and E18.5 (**B**). % oocytes in pre-leptotene, leptotene, zygotene, pachytene, and diplotene stages was determined by synaptonemal complex morphology visualized by SYCP3 immunofluorescence labeling on meiotic spreads. Statistical significance was determined by Chi square test. ns p>0.05, **p<0.01 (table S1).

**Fig. S3.**
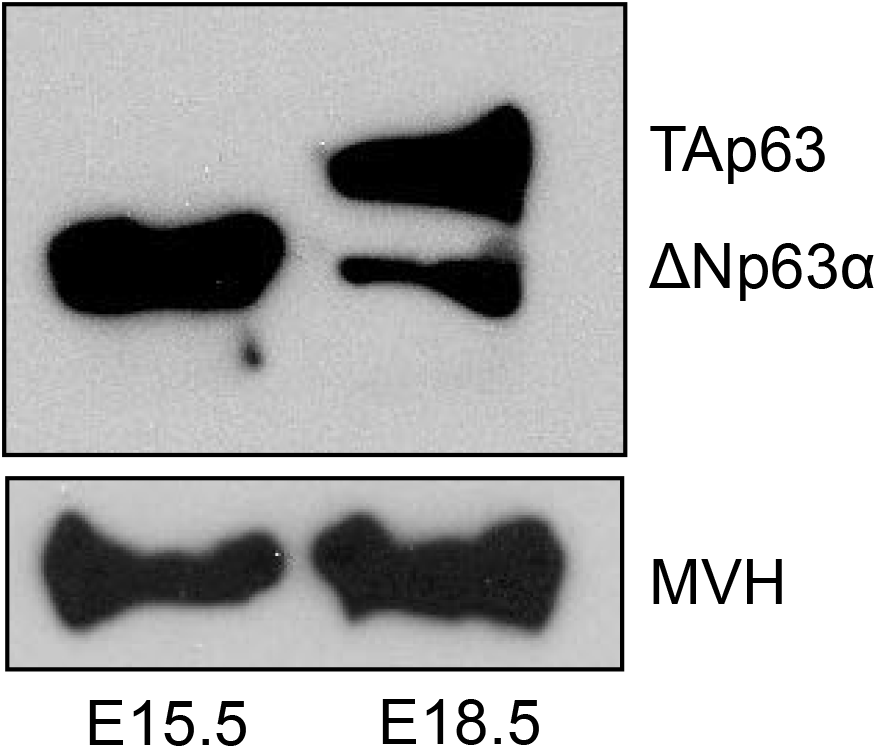
Expression of p63 isoforms in early and late MPI. Western blot detection of p63 dominant negative (ΔNp63) and transactivating (TAp63) isoforms in E15.5 and E18.5 ovary lysates of B6 wild-type mice. Western blot detection of mouse vasa homolog (MVH) in same lysates used as a loading control.

**Fig. S4.**
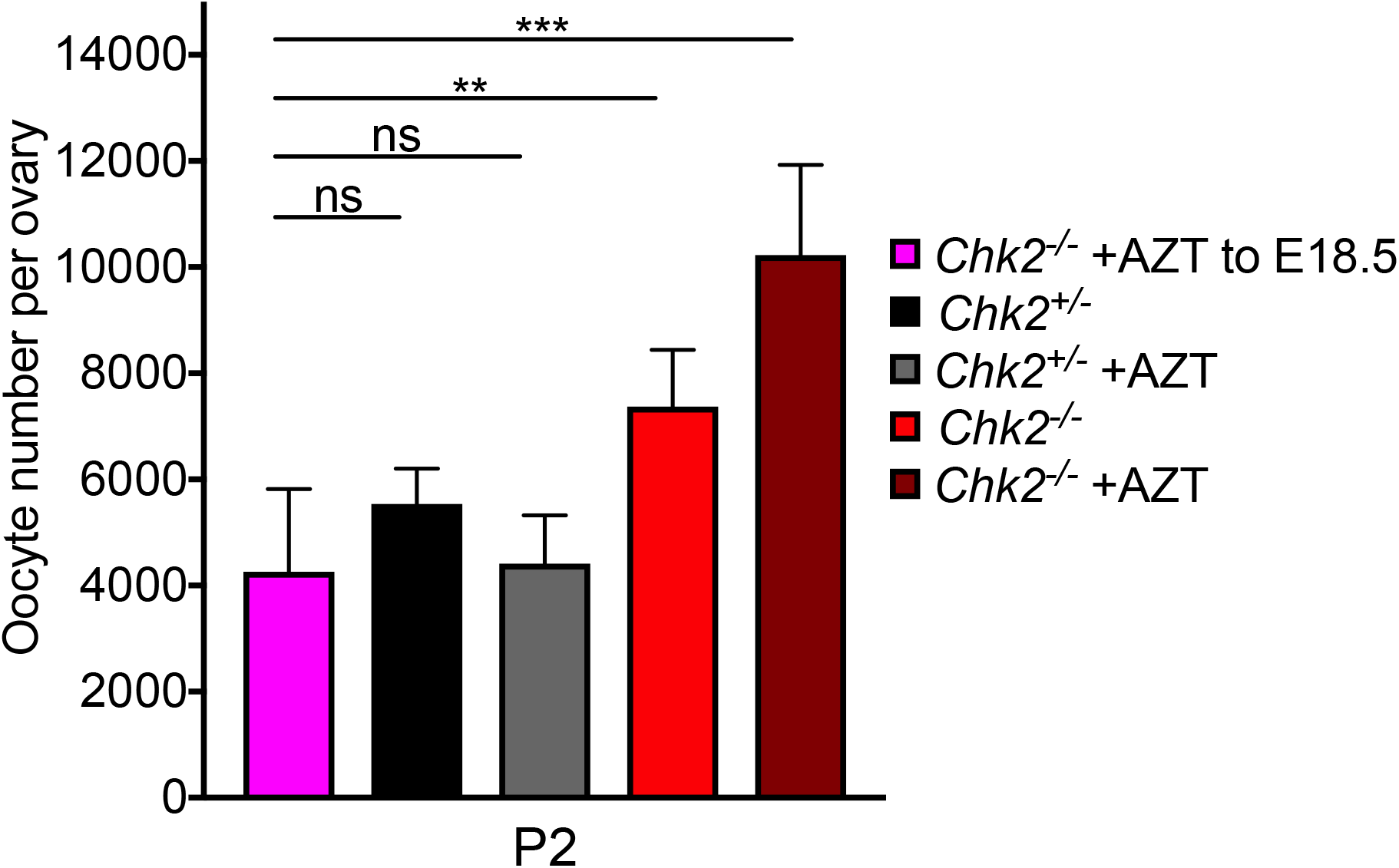
Mechanisms driving FOA are not temporally distinct. Quantification of oocytes per ovary in P2 *Chk2*^−/−^ mice treated with AZT until E18.5, followed by no treatment until P2. Compared to oocyte number per ovary at P2 in *Chk2*^+/−^ and *Chk2*^−/−^ untreated and AZT-treated results from Fig. 1E, table S2. Statistical significance was determined using unpaired t-test, ns p>0.05; **p<0.01; ***p<0.001.

**Fig. S5.**
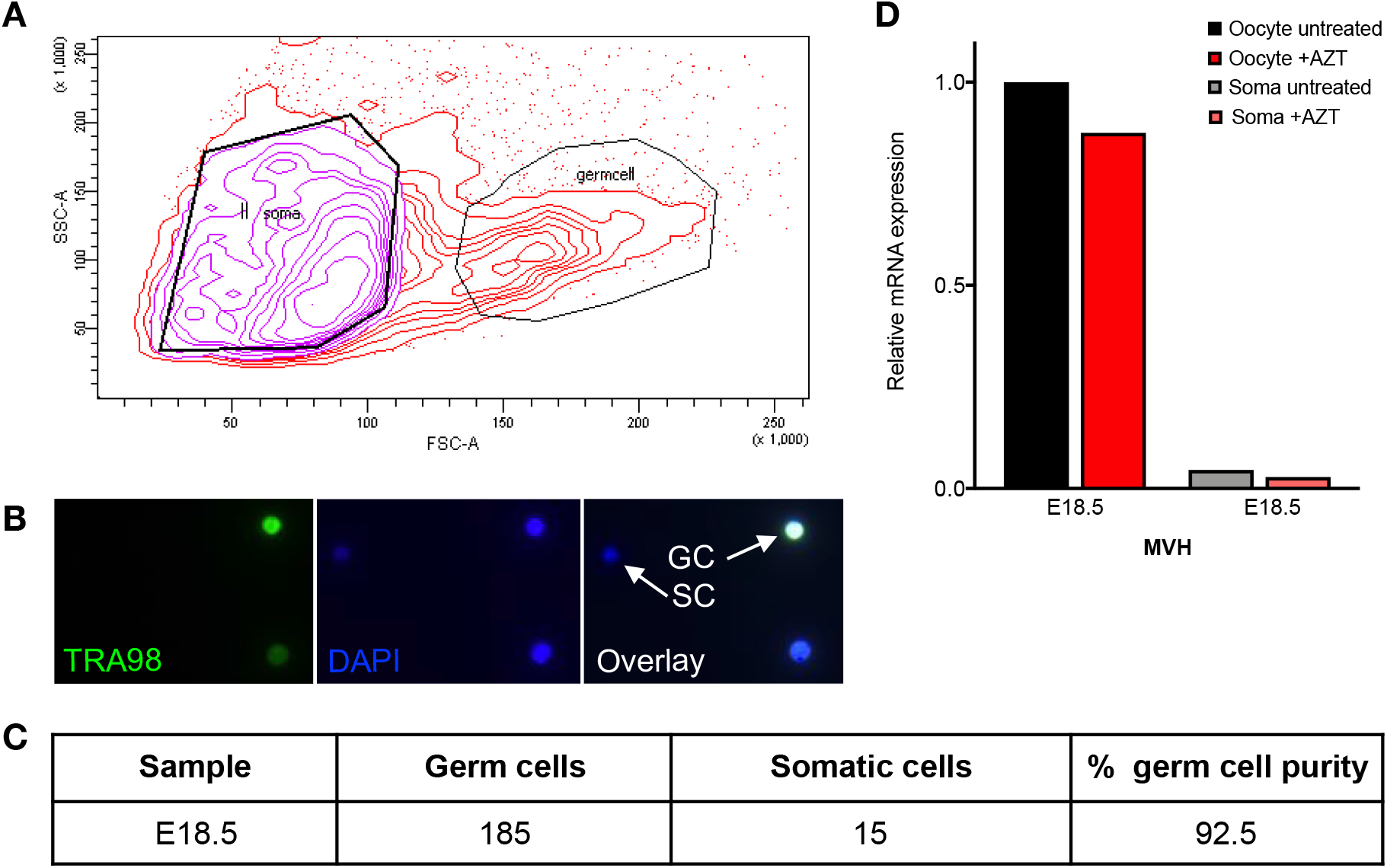
Oocyte FACS-sorting method and purity assessment. (**A**) FACS profile during isolation of oocytes from ovarian somatic cells based on side and forward scatter parameters. (**B**) Analysis of oocyte purity in sorted oocyte sample based on TRA98 and DAPI staining. (**C**) Quantification of oocytes vs somatic cells in sorted oocyte sample with percent oocyte purity. (**D**) Analysis of oocyte purity in untreated and AZT-treated sorted oocytes and ovarian somatic cells using quantitative RT-PCR detection of germ cell-specific gene *Mvh*. Relative expression normalized to that of the untreated oocyte sample using the *Actb* gene as an internal control. All mice used were of CD1 genetic background. Each sample contained oocytes or ovarian somatic cells from at least 6 embryos.

**Fig. S6.**
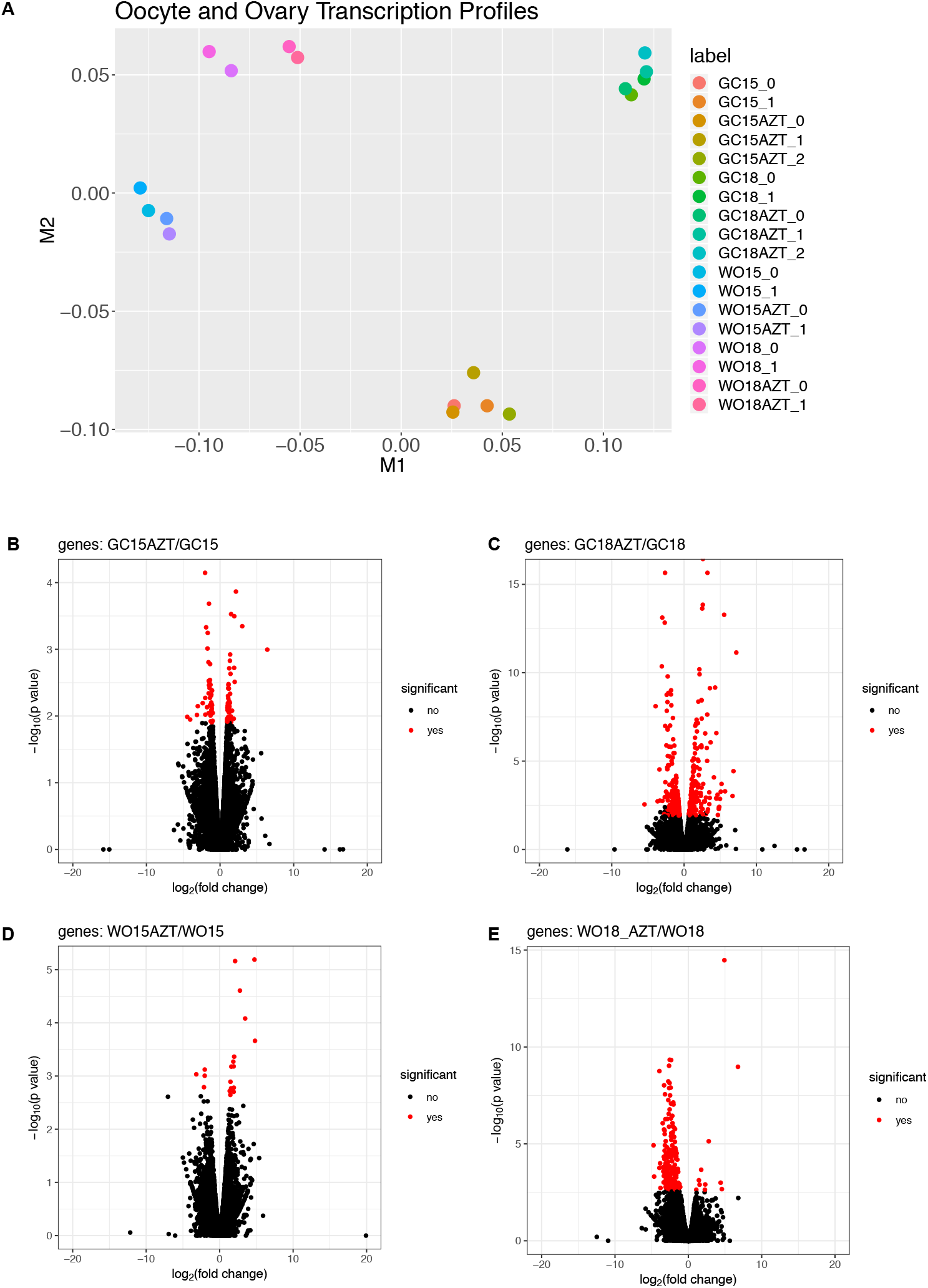
Gene expression profiles from bulk RNA-Seq samples. (**A**) Multidimensional scaling (MDS) plot to visualize the similarities in global gene expression between whole ovary (WO) and germ cell/oocyte (GC) samples and biological replicates (**B-E**) Volcano plots displaying pairwise comparison of differential gene expression between E15.5 untreated and E15.5 +AZT oocytes (**B**), E18.5 untreated and E18.5 +AZT oocytes (**C**), E15.5 untreated and E15.5 +AZT ovaries (**D**), and E18.5 untreated and E18.5 +AZT ovaries (**E**) (table S3A-D). Y-axis range of volcano plots varies between pairwise comparisons. All mice used were of CD1 genetic background.

**Fig. S7.**
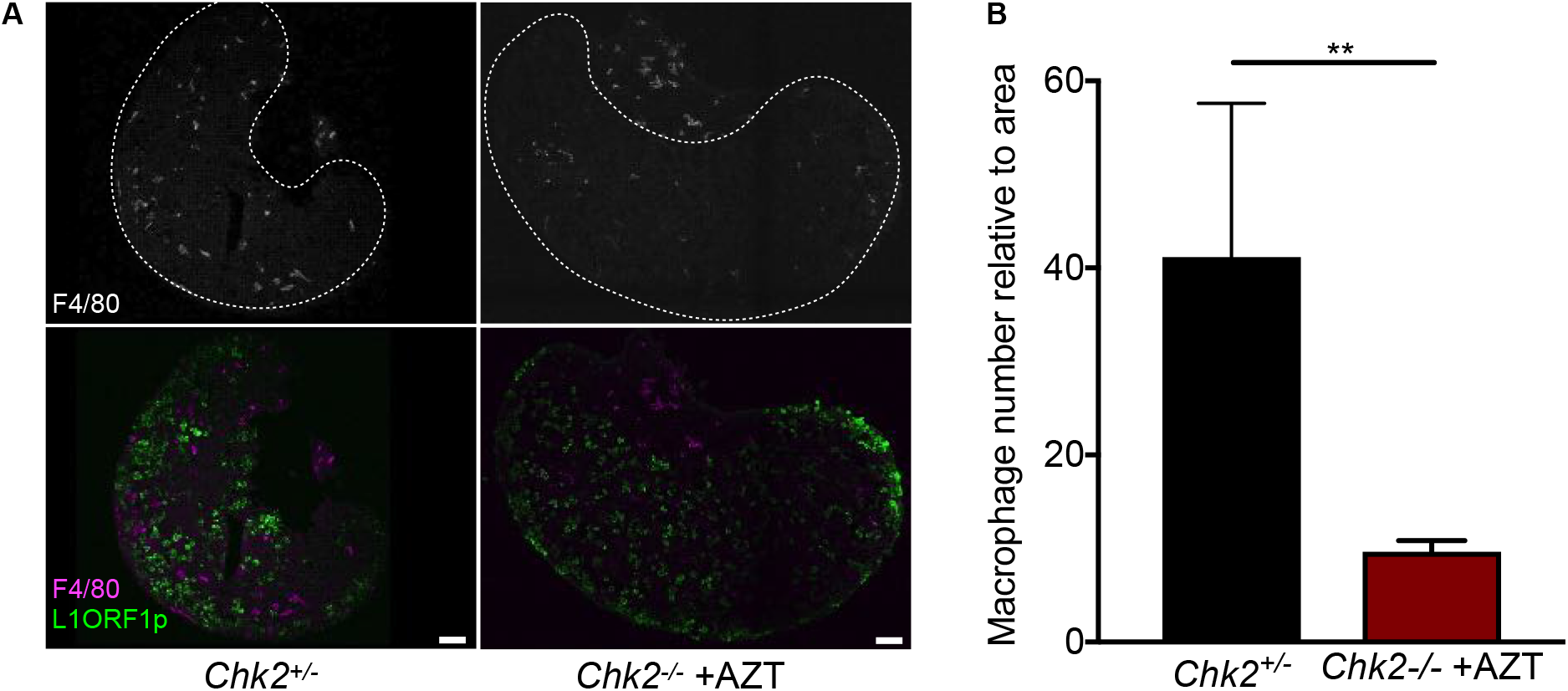
Macrophage occupation of ovary in untreated and AZT-treated conditions. (**A**) Immunofluorescence labeling with macrophage marker F4/80 and L1 ORF1p to label oocytes in E18.5 ovaries of *Chk2*^+/−^ and *Chk2*^−/−^ +AZT conditions. Ovary separated from soma with dotted white line. Scalebar represents 50μm. (**B**) Quantification of macrophage number per ovary area in *Chk2*^+/−^ and *Chk2*^−/−^ +AZT conditions. Statistical significance determined using Mann-Whitney test, p**<0.01.

**Fig. S8.**
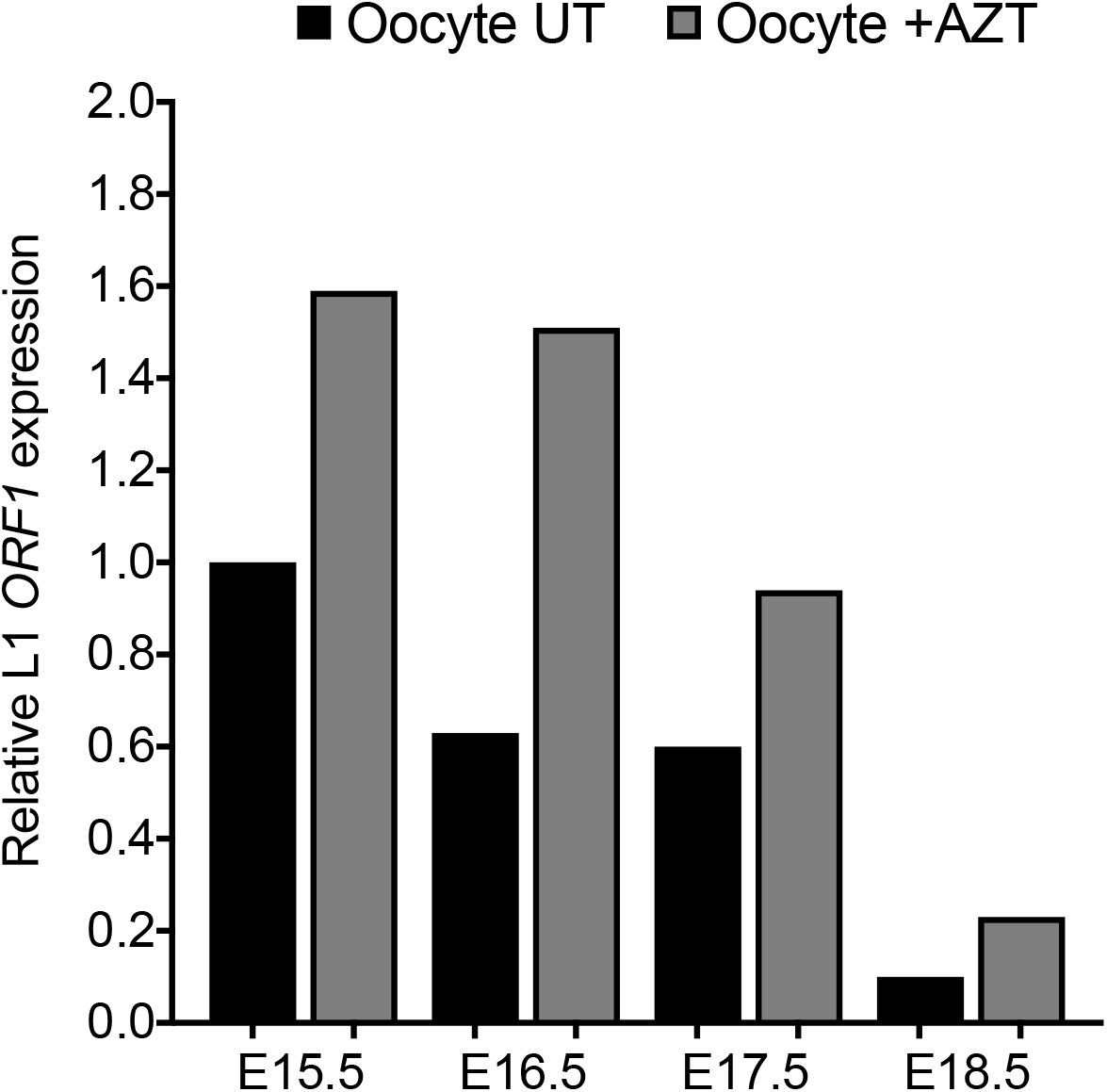
L1 *ORF1* mRNA expression in untreated and AZT-treated oocytes. Quantitative RT-PCR detection of L1 *ORF1* relative mRNA expression at E15.5, E16.5, E17.5, and E18.5 in untreated and AZT-treated FACS-sorted oocytes. Expression normalized to that of the E15.5 untreated oocyte sample using the *Actb* gene as an internal control. All mice used were of CD1 genetic background. Each sample contained oocytes from at least 6 embryos.

**Fig. S9.**
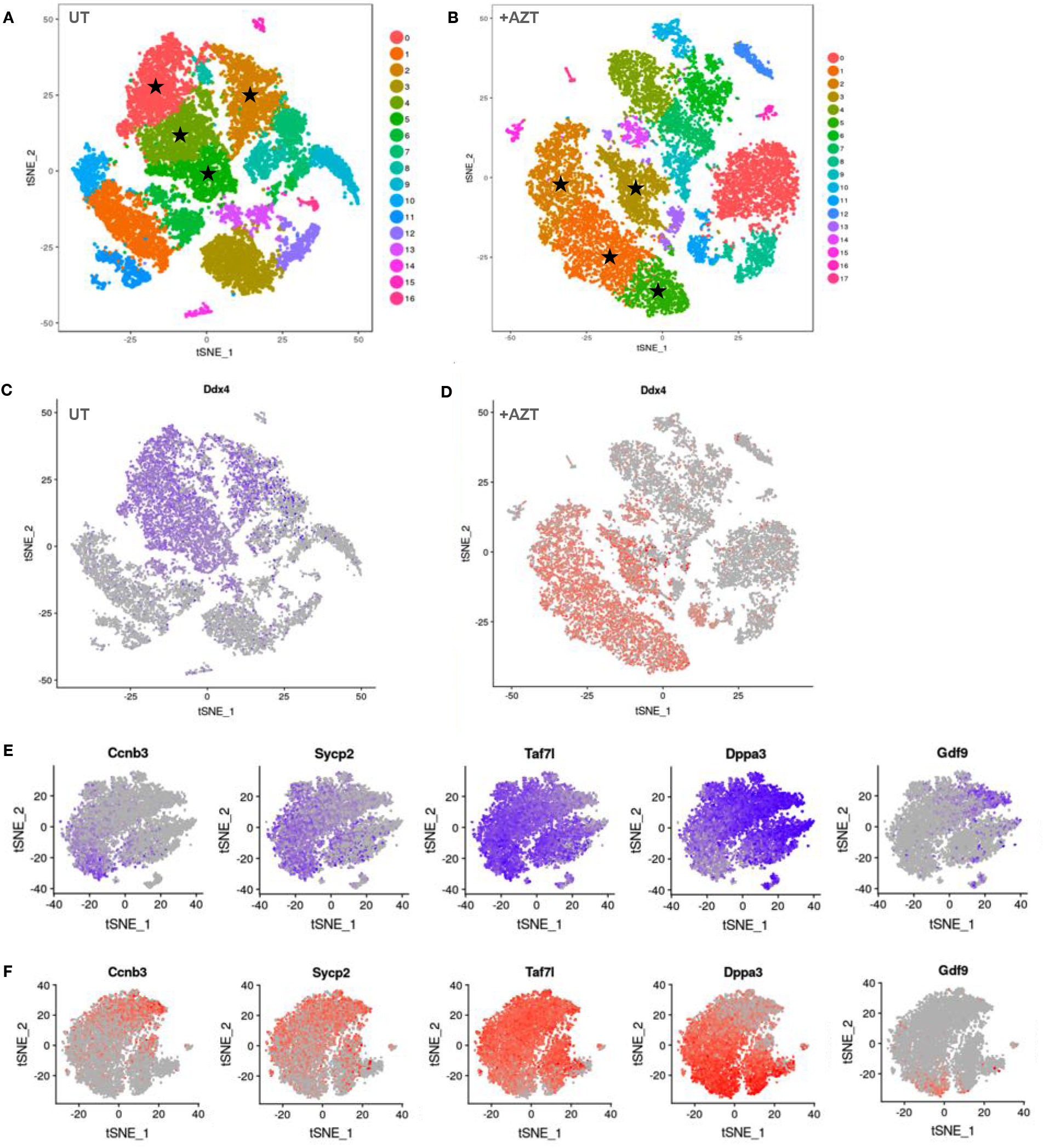
Identification of early and advanced oocyte clusters in single-cell RNA-Seq. (**A**) t-SNE plot of untreated ovarian cell clusters. (**B**) t-SNE plot of AZT-treated ovarian cell clusters. Black stars indicate clusters expressing germ cell-specific gene *Ddx4* that were used for re-clustering of oocytes. (**C-D**) t-SNE plot highlighting germ cell-specific gene *Ddx4* in blue in untreated (**C**) and red in AZT-treated (**D**) ovarian cells. (**E-F**) t-SNE plots showing germ cell-specific genes ranging from early (Ccnb3), intermediate (Sycp2, Taf71, Dppa3), and late (Gdf9) developmental markers in untreated (**E**) and AZT-treated (**F**) re-clustered oocyte populations. All embryos used were at E18.5 and of CD1 genetic background.

**Table S1. Quantification and statistical analysis for meiotic progression in *Chk2*+/− and *Chk2*−/− oocytes.**

**Table S2. Quantification and statistical analysis for *Chk2*+/− and *Chk2*−/− untreated and AZT-treated ovary experiments.**

**Table S3. Differential gene expression and gene ontology for untreated and AZT-treated oocytes and ovaries.**

**Table S4. Differential gene expression in untreated and AZT-treated single-cell clusters.**

**Table S5. Small RNA length distribution and repeat alignment.**

**Table S6. Quantification and statistical analysis for *Mili;Chk2* untreated and AZT-treated ovary experiments.**

**Table S7. Quantification of follicles and fertility in *Chk2*+/− untreated and *Chk2*−/− untreated and AZT-treated mice.**

